# Consonance perception in congenital amusia: behavioral and brain responses to harmonicity and beating cues

**DOI:** 10.1101/2021.11.15.468620

**Authors:** Jackson E. Graves, Agathe Pralus, Lesly Fornoni, Andrew J. Oxenham, Barbara Tillmann, Anne Caclin

## Abstract

Congenital amusia is a neurodevelopmental disorder characterized by difficulties in the perception and production of music, including the perception of consonance and dissonance, or the judgment of certain combinations of pitches as more pleasant than others. Two perceptual cues for dissonance are inharmonicity (the lack of a common fundamental frequency between components) and beating (amplitude fluctuations produced by close, interacting frequency components). In the presence of inharmonicities or beats, amusics have previously been reported to be insensitive to inharmonicity, but to exhibit normal sensitivity to beats. In the present study, we measured adaptive discrimination thresholds in amusic participants and found elevated thresholds for both cues. We recorded EEG and measured the mismatch negativity (MMN) in evoked potentials to consonance and dissonance deviants in an oddball paradigm. The amplitude of the MMN response was similar overall for amusics and controls, but while control participants showed a stronger MMN to harmonicity cues than to beating cues, amusic participants showed a stronger MMN to beating cues than to harmonicity cues. These findings suggest that initial encoding of consonance cues may be intact in amusia despite impaired behavioral performance, but that the relative weight of non-spectral cues may be increased for amusic individuals.

## 1. INTRODUCTION

A fundamental aspect of Western tonal music is the phenomenon of consonance and dissonance, whereby certain combinations of pitches are judged to be more pleasant, and to create less tension, than others. The perception of consonance is likely to be determined at least in part by enculturation (Butler & Daston, 1968; Lundin, 1947; McDermott, Schultz, Undurraga, & Godoy, 2016), but some aspects of consonance may arise from the acoustic properties of musical sounds. Pitch intervals with simple frequency ratios are often rated as more consonant than complex ratios, a phenomenon with at least two potential explanations. Dissonant (i.e., complex ratio) intervals generally produce *beating*: upper harmonics of the two tones are close together in frequency and interact in the auditory periphery, leading to amplitude fluctuations that are perceived as roughness, which is often described as unpleasant (Hutchinson & Knopoff, 1978; Plomp & Levelt, 1965). Dissonant intervals also lack *harmonicity*: their components cannot be interpreted as multiples of a common fundamental frequency (Ebeling, 2008; Terhardt, 1974). One study testing the relative importance of these two cues found that inharmonicity was likelier than beating to underlie the perception of dissonance (McDermott et al., 2010).

Further insight into consonance perception comes from the study of congenital amusia, a disorder characterized by difficulties in the perception and production of music in individuals with clinically normal hearing and no general cognitive deficits (Ayotte et al., 2002), which affects 2-4% of the population (Kalmus & Fry, 1980; Peretz & Vuvan, 2017). Deficits exhibited by individuals with amusia can provide insights about general mechanisms of music perception, and of auditory perception, more generally. Pleasantness ratings of chords have been found to be impaired to some degree in congenital amusia (Cousineau et al., 2012; Marin et al., 2015). One of these previous studies tested perception of consonance and dissonance cues in amusics and found that they exhibit behavioral impairment for the detection of inharmonicity, but not for the detection of beating (Cousineau et al., 2012), lending further support to the view that consonance perception is dominated by harmonicity (McDermott et al., 2010).

Congenital amusia is generally thought of as a disorder affecting pitch perception (Cousineau et al., 2015; Peretz, 2016; Tillmann et al., 2015) and timbral cues relating to the frequency content of sounds (Graves et al., 2019; Marin et al., 2012), leaving intact other capabilities, such as memory for speech stimuli (Albouy, Peretz, et al., 2019; Tillmann et al., 2009), and with more restricted or conditional deficits for other musical attributes, like rhythm (Foxton et al., 2006; Hyde & Peretz, 2004) and loudness (Graves et al., 2019; Tillmann et al., 2016). However, a recent study (Whiteford & Oxenham, 2017) found that amusics are impaired at detecting amplitude modulation (AM) as well as frequency modulation (FM), suggesting that the deficit may not be limited to cues related to pitch and timbre. The finding that amusics are impaired at detection of AM has interesting implications for amusic perception of beating, and seems to conflict with the previously reported lack of amusic impairment for beating perception (Cousineau et al., 2012). This discrepancy may be due to the different levels of AM tested by the two studies: Cousineau et al. (2012) used stimuli that produced the equivalent of 100% depth of AM (well above threshold), whereas Whiteford and Oxenham (2017) measured AM detection at modulation depths near threshold. However, thresholds for detection of beating in a complex tone have not been directly measured for amusics, and further exploration of consonance perception in amusia is warranted to resolve this apparent discrepancy.

Studies of brain function and anatomy in amusia generally point to an origin of the deficit in the network connecting auditory areas in temporal cortex with higher-level areas in frontal cortex (Albouy et al., 2013, 2015; Hyde et al., 2006, 2007, 2011; Leveque et al., 2016). In terms of early cortical encoding of pitch, fMRI studies did not observe differences in amusics’ activation patterns with classical subtraction analyses (Hyde et al., 2011; Norman-Haignere et al., 2016), but 1) a re-analysis with whole-brain multivariate pattern analyses revealed that amusics and controls differed in their pattern of functional activity in right Heschl’s gyrus (Albouy, Caclin, et al., 2019) and 2) an MEG study observed differences in amusics and controls at this early level (Albouy et al., 2013), leaving open the possibility that early pitch processing may also be impacted in amusia.

Automatic sound processing in amusics has previously been studied using the mismatch negativity (MMN), an EEG potential recorded in response to sounds that violate low-level auditory expectations, and that can occur even in the absence of directed attention (Näätänen et al., 1978, 2007). Despite showing behavioral impairment at pitch discrimination tasks, amusics have been found to exhibit normal MMN responses even to small pitch changes that are below their behavioral discrimination threshold (Moreau et al., 2009, 2013; Peretz et al., 2009). A similar pattern of overall normal MMNs in amusia has also recently been observed for emotional prosody (Pralus et al., 2020) and in pitch sequences of various levels of melodic complexity (Quiroga-Martinez, Tillmann, et al., 2021), yet with differences in an early negativity preceding the MMN and in the P3a (Pralus et al., 2020). One EEG study found a reduced early frontal negativity in response to unpredictable notes in a melody (Omigie et al., 2013), suggesting that potential impairments of in automatic brain responses at this early level may still exist for amusics for certain perceptual cues.

For normal-hearing, non-amusic listeners, the MMN has also been reported in response to harmonicity changes and to pitch differences of harmonic complex tones (Butler & Trainor, 2012; Tervaniemi, Schröger, Saher, & Näätänen, 2000). In addition to high-level cues like tonal function (Marmel et al., 2011), the MMN has been observed in response to differences in consonance and dissonance (Brattico et al., 2009; Crespo-Bojorque et al., 2018; Linnavalli et al., 2020). More specifically, an MMN has been measured in response to inharmonicity in the form of a mistuned harmonic (Alain et al., 2001) and to harmonicity in the form of a harmonic complex presented in the context of inharmonic complexes (Jones, 2003). The MMN thus makes a good candidate for indexing early neural processing of consonance and dissonance in amusics, as its presence has been established in response to pitch changes for amusics, and in response to dissonance cues for normal-hearing listeners.

In the present study, we tested whether behavioral sensitivity and MMN differed for amusics and controls in response to two potential cues for consonance and dissonance: harmonicity and beating. First, participants’ detection thresholds were measured for both cues as well as for pure-tone AM. If perception of beating depends on AM detection, any impairment for pure-tone AM detection (Whiteford & Oxenham, 2017) should be reflected in elevated thresholds for detection of beating in a complex tone. Second, in an EEG study, participants listened to streams of stimuli in an oddball paradigm, where consonant or dissonant stimuli deviated from a standard stimulus. If behavioral impairments for consonance perception can be traced to differences in early encoding of these cues, this could be reflected in reduced MMN amplitude for amusics. If on the other hand the deficits arise from later processes, amusics should show normal MMN in response to these stimuli.

## 2. METHODS

### 2.1 Participants

For the study measuring behavioral thresholds, we recruited 12 amusic individuals (6 female and 6 male) and 11 control participants (7 female and 4 male). For the EEG study, we recruited 19 amusic individuals (10 female and 9 male) and 21 control participants (13 female and 8 male), of which 13 individuals (5 amusics and 8 controls) had also participated in the behavioral study. Participants in the EEG study took part in two consecutive MMN studies within the same session; the first one with emotional vowels is reported in Pralus et al. (2020) and the second one is reported here.

During recruitment, control participants were matched to amusic participants in terms of age, education, gender, and handedness (see Table 1). No participants had any formal musical training, except for two control participants who each reported one year of musical training (one participated only in the behavioral study and the other only in the EEG study). All participants had normal hearing as determined by pure tone audiometry, and congenital amusia was identified with the Montreal Battery of Evaluation of Amusia (MBEA) (Peretz et al., 2003). All participants provided written informed consent and were compensated for their participation. Study procedures were approved by the required ethics committee, the Comité de Protection des Personnes (CPP), Sud-Est II.

**Table 1.**
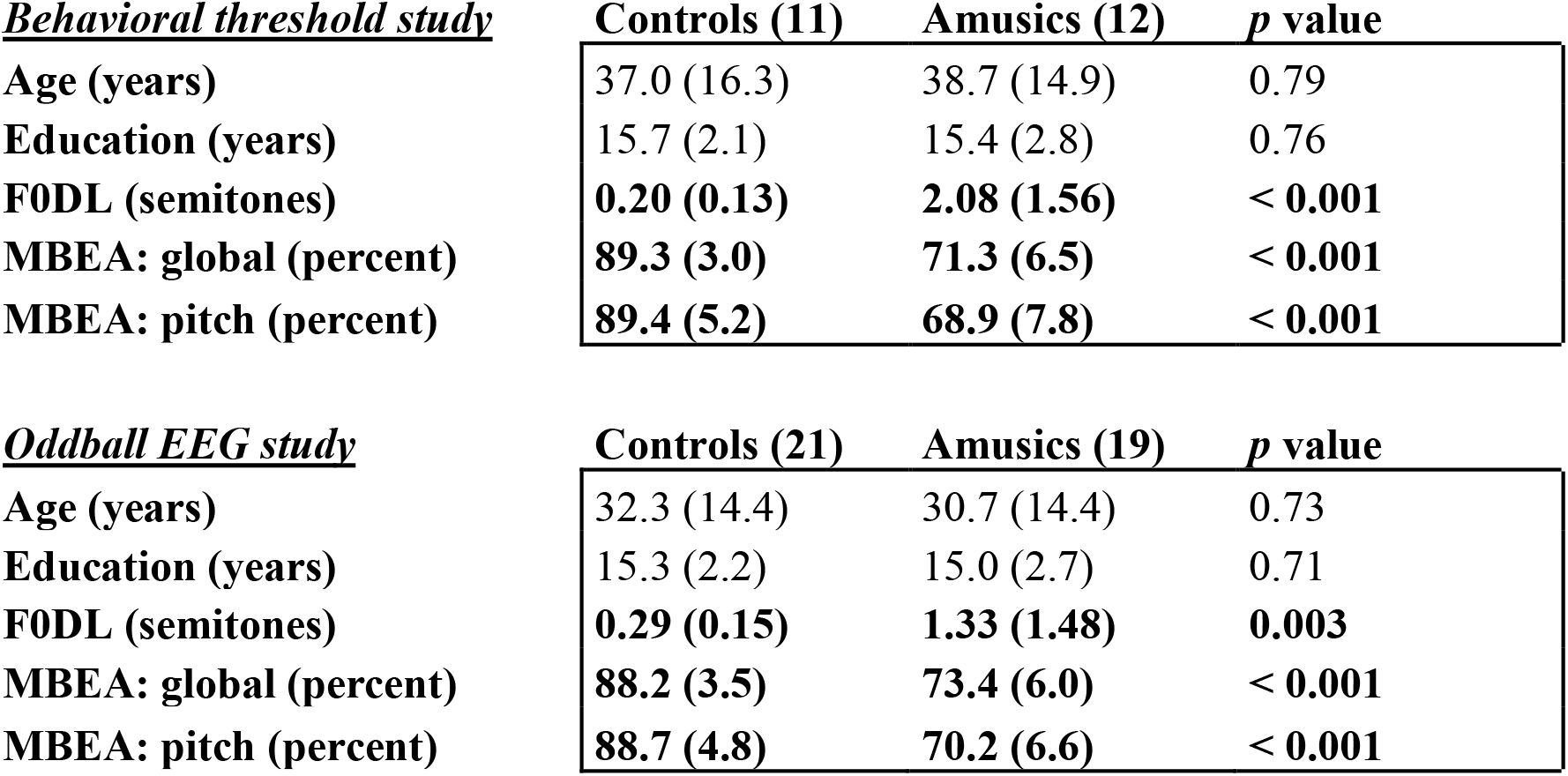
Comparison of amusic and control samples by demographic variables and music perception scores. Mean values are listed, with standard deviations in parentheses. Bold indicates a *p* value less than 0.05 in a two-tailed independent t-test between groups. Congenital amusia is identified using the Montreal Battery of Evaluation of Amusia (MBEA), with a global cutoff score of 78% correct over six subtests (Peretz et al., 2003). MBEA scores for the pitch-related subtests only (Scale, Interval, and Contour) are also listed, along with fundamental frequency difference limens (F0DL) measured using an adaptive tracking procedure (Tillmann et al. 2009). Thirteen participants completed both studies, along with 10 completing only the behavioral study and 27 completing only the EEG study. All participants except two control participants (one in each study, see main text) reported 0 years of musical experience.

### 2.2 Stimuli

Complex tones, partly inspired by stimuli used by Cousineau et al. (2012), were generated in four categories in order to manipulate consonance cues: harmonic, inharmonic, no-beating, and beating (see Figure 1). Harmonic complexes consisted of the 1^st^, 2^nd^, 3^rd^, 5^th^, and 9^th^ components of a harmonic series, added together in sine phase. Inharmonic complexes were generated by shifting every component in a harmonic complex up by a uniform distance in linear frequency. When manipulating beating cues, we wanted to be sure that the perception of dissonance was not due to inharmonicity from the introduction of non-harmonic components, but purely due to beating. Therefore, no-beating stimuli were always inharmonic, with frequency components chosen by sampling from a uniform distribution between −30 and +30 Hz relative to each component in the equivalent harmonic stimulus, added together in random phase. Beating stimuli were created by adding a single sideband component 30 Hz away from each component in a no-beating stimulus, with a uniform level difference between sidebands and original components. The location of sidebands (above or below original components) was randomly chosen for each complex, but consistent across the complex. See supplemental audio files SA1 - SA4 for examples of complexes in each of the four categories.

**Figure 1.**
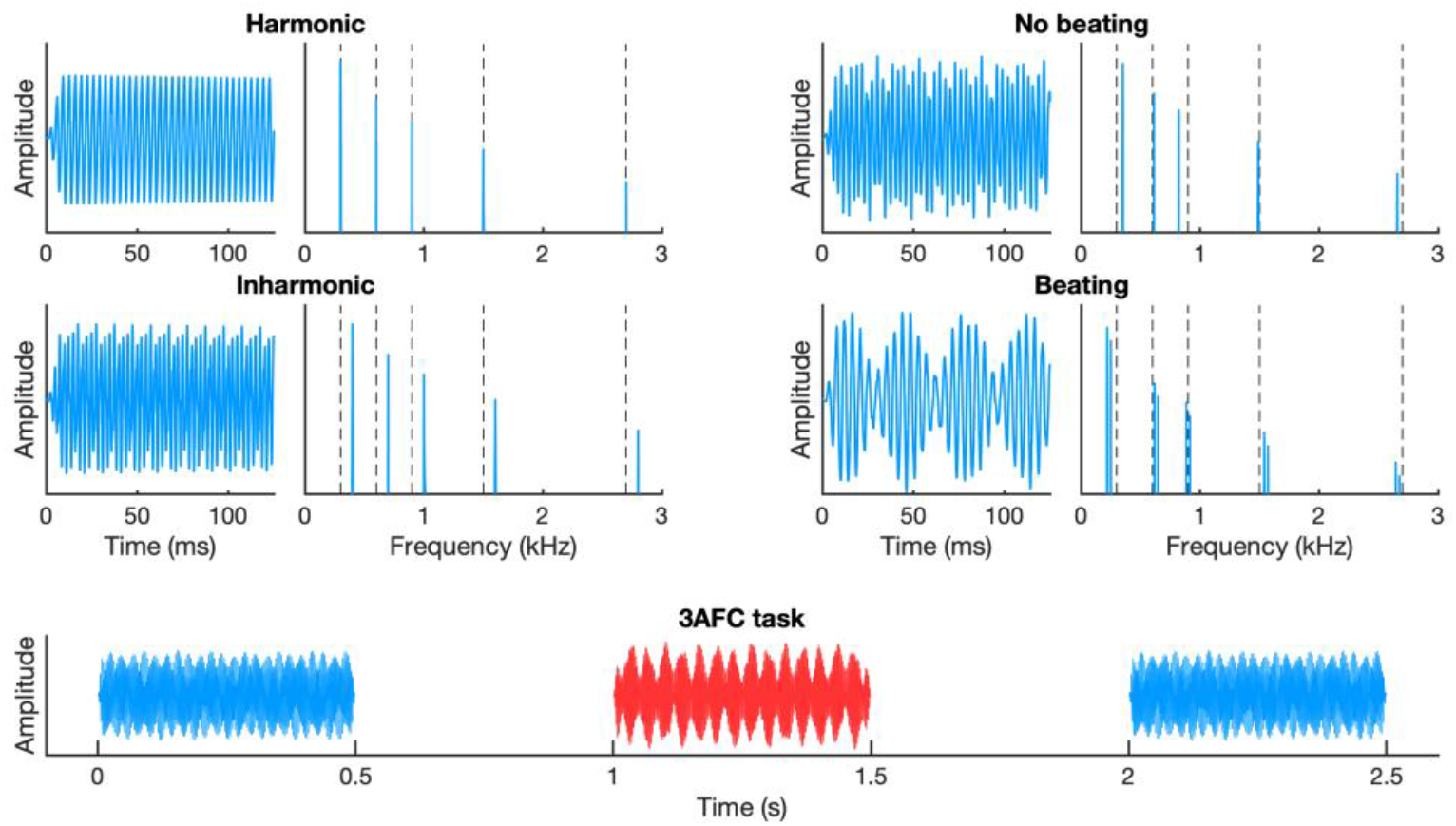
Stimuli manipulating harmonicity and beating cues. Temporal waveforms and frequency spectra of stimuli for harmonicity (left) and beating (right) conditions, with vertical dashed lines showing the 1^st^, 2^nd^, 3^rd^, 5^th^, and 9^th^ components in a harmonic series. **Left**: harmonic complexes contained the five components shown, with a decaying spectral envelope. Inharmonic complexes were shifted up relative to harmonic complexes by a uniform distance in linear frequency. **Right:** no-beating stimuli were inharmonic, with frequency components jittered relative to a harmonic series. Beating stimuli introduced a single sideband component 30 Hz above or below each component in the no-beating stimuli, with a uniform level difference between sidebands and original components. **Bottom**: timing of a trial on the 3AFC discrimination task used for threshold measurement. Participants identified the target (red) by comparing against two references (blue), with target position randomized on each trial.

In addition to our manipulation of beating, which indirectly introduced AM to a complex tone, we also directly measured thresholds for detection of pure-tone AM. Stimuli for the AM detection test involved a pure-tone carrier with a frequency that was roved between 277 and 370 Hz. When AM was present, the modulation frequency was constant at 10 Hz, and the depth was adaptively varied to measure threshold.

In order to ensure that perception of dissonance cues was based on the dissonance itself and not on a perceived difference in pitch, we varied the fundamental frequency (F0) between intervals of all tones used in both studies. The F0 of each complex tone (or carrier frequency for AM stimuli, or nominal F0 for inharmonic, no-beating, and beating stimuli) was chosen from among six discrete values, at semitone increments between C#4 (277 Hz) and F#4 (370 Hz). Throughout the behavioral threshold study and the oddball MMN study, no two consecutive tones shared an F0. All complex tone stimuli had a decaying spectral envelope, such that components decreased in level at a rate of 14 dB/octave from the lowest component. The duration of each tone was 500 ms, with 10-ms Hanning window on- and off-ramps, and the overall level of each tone was 65 dB SPL. All stimuli were created using MATLAB (The Mathworks, Natick, MA) and presented at a sampling rate of 48 kHz through over-ear headphones in a quiet sound booth.

For the behavioral threshold study, the size of the spectral shift for inharmonic stimuli and the amplitudes of the sidebands for beating stimuli were adapted in order to measure a threshold. During these adaptive procedures, the size of the inharmonic frequency shift varied on a logarithmic scale between 1 and 125 Hz and the amplitude of the sidebands varied between −60 and 0 dB relative to the original components. For the oddball MMN study, a constant shift size and sideband amplitude were chosen to differentiate the standard and deviant stimuli to be played to every participant. These levels were chosen in order to maximize the difference in discriminability between participant groups (amusic and control). Accordingly, we chose an inharmonic shift of 12 Hz, which was higher than 82% of controls’ individual thresholds but only 42% of amusics’ individual thresholds, and a sideband amplitude of −19 dB (relative to the original tones), which was higher than 82% of controls’ individual thresholds but only 33% of amusics’ individual thresholds.

### 2.3 Procedure

#### 2.3.1. Measurement of behavioral discrimination thresholds

In the behavioral threshold study, participants discriminated inharmonic from harmonic, beating from no-beating, or AM from no-AM stimuli in a 3AFC task. On each trial, participants heard three tones, and were asked to select the tone that sounded different from the other two. The target tone was always the more dissonant sound (inharmonic, beating, or AM), while the two reference tones were more consonant (harmonic, no beating, or no AM). Tones had durations of 500 ms, with 500 ms silent gaps in between tones. The three tones always had three different F0s: C#4, D#4, and F4, or D4, E4, and F#4, presented in a random order that was independent of which tone was the target. Listeners were thus carefully instructed to ignore changes in F0 and identify the tone that stood out from the other two (without explaining explicitly on which feature).

The amount of difference between target and reference was adaptively varied, and discrimination thresholds were measured using a tracking procedure following a 1-up, 2-down rule. An adaptive run began at 50 Hz and changed by factors of 2.51, 1.58, and 1.26 (inharmonic shift) or began at −10 dB and changed by step sizes of 4, 2, and 1 dB (sideband and AM depth). Step size changed every 2 reversals until the final step size, after which the run continued until 6 reversals were completed at the final step size. Each participant completed three runs for each cue (harmonicity, beating, and AM) and the averages of the six last reversals from each run (geometric averages for inharmonicity) were averaged together for an estimated threshold.

#### 2.3.2. MMN oddball paradigm

During EEG recording, streams of tones were presented to participants with a stimulus onset asynchrony (SOA) of 700 ms. In each of four conditions, one stimulus type was defined as standard, occurring on 5/6 of trials, and its opposite as deviant, occurring on only 1/6 of trials (see Figure 2). Conditions were named after the deviant tone: harmonic-deviant (HDEV), inharmonic-deviant (IDEV), no-beating-deviant (NDEV), and beating-deviant (BDEV). Deviants were always separated from each other in the stream by no more than 7 and no fewer than 3 standards. The F0 of each tone was selected randomly, with the constraint that no two consecutive tones shared an F0. Each participant completed four blocks of passive listening followed by four blocks of an active task (see below). Passive blocks always came before active blocks. The order of the four blocks was always the same in passive and active phases, but the order of blocks was counterbalanced across participants. Passive blocks contained 140 deviants and 700 standards in total, and participants were instructed to sit still and watch a silent subtitled movie. Active blocks contained 30 deviants and 150 standards, and participants were instructed to press a button every time they heard a sound that was different from most other sounds.

**Figure 2.**
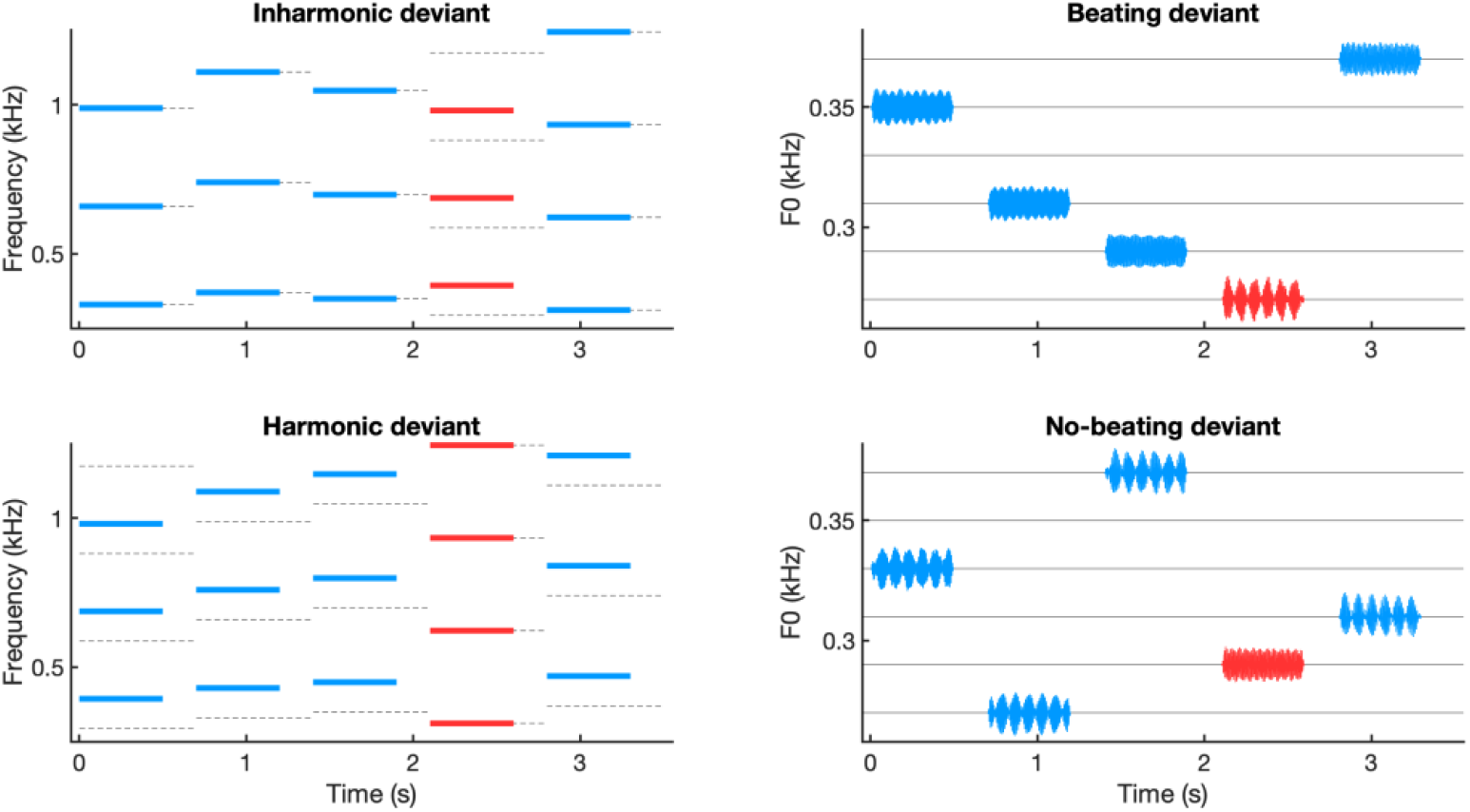
Oddball paradigm with consonance changes and pitch roving. Participants heard deviant (red) and standard (blue) stimuli in four different experimental conditions, while nominal F0 varied constantly. Each of the four stimulus types (harmonic, inharmonic, no-beating and beating) served as the deviant in one condition, with its contrast serving as the standard. Left panels: sound spectra for inharmonic-deviant and harmonic-deviant conditions, dashed grey lines indicate harmonic partials. Right panels: sound waveforms for beating-deviant and no-beating-deviant conditions, plotted at the level of their fundamental frequency (F0). In the beating-deviant and no-beating-deviant conditions, all sounds were inharmonic.

As a measure of behavioral sensitivity to the difference between standard and deviant, the sensitivity index d′ was calculated from active-task responses in the following manner. Any responses between 100 and 2200 ms after a deviant were counted as a hit for that deviant, and any responses between 2200 ms after a deviant and 100 ms after the following deviant were counted as a false alarm. In this way, the duration of hit periods was constant at 2100 ms per deviant, while the duration of false alarm periods was variable from 700 to 3500 ms, but averaged 2100 ms, meaning that a random responder would make an equal number of hits and false alarms. Sensitivity d′ was then calculated using the resulting hit and false alarm rates (Stanislaw & Todorov, 1999).

### 2.4. EEG recording and preprocessing

EEG was recorded from 31 Ag/AgCl active electrodes (Acticap, Brain Products GmbH, Gilching, Germany), with a nose reference, at a sampling rate of 1000 Hz with a 0.016-1000 Hz bandwidth (BrainAmp Standard Amplifier, Brain Products GmbH, Gilching, Germany). Vertical eye movements were recorded with an additional active electrode positioned under the left eye (offline re-referenced to Fp1). Pre-processing was conducted in MATLAB using functions from eeglab (Delorme & Makeig, 2004). Data was bandstop filtered in order to remove line noise, using 4^th^-order Butterworth filters with cutoffs of 47 and 53 Hz, then 147 and 153 Hz. A 20-second-long portion of data containing eyeblinks but few other artifacts was visually selected for each participant, for the purpose of performing an independent components analysis (ICA). ICA weights from this subset of data were visually examined, eyeblink components were selected and then removed from the entirety of the data through an ICA inverse transformation. After ICA-based blink correction, the data was divided into 700-ms epochs, from −200 ms to +500 ms relative to stimulus onset, and non-blink artifacts were rejected using a dynamic range threshold applied to each epoch. This threshold, and the channels to be ignored for artifact rejection, were manually selected for each participant based on visual inspection of the data (thresholds ranged from 90 μV to 300 μV, with an average of 153 μV; the number of excluded channels ranged from 0 to 21, with an average of 7.59). Of the 140 deviant trials in each passive block, the average number rejected due to artifacts was 31, leaving an average of 109 deviants for analysis per passive block. After artifact rejection, epochs were averaged for each condition, separately for deviants and standards, excluding standards immediately following a deviant. The resulting ERPs were bandpass filtered (Butterworth, 4^th^ order) from 2 to 30 Hz, and the baseline from 100 ms to 0ms before stimulus onset was subtracted. Finally, any exceptionally noisy electrodes were removed and re-interpolated from surrounding electrodes based on their three-dimensional coordinates.

### 2.5. Analysis of EEG data

Only EEG data from the passive listening blocks were analyzed. The MMN was evaluated by subtracting ERPs to standard sounds from their acoustically identical deviant sounds. This means, for example, that the harmonic deviant (from the HDEV block) was compared against the harmonic standard (from the IDEV block), not against the (inharmonic) standard from its own block. In this way, any measured difference between ERPs cannot be due to acoustic differences between the sounds, as they are the same sound, and is likely instead to arise from neural encoding of the deviant sound as deviant. For each deviant-standard pair, ERP amplitudes were evaluated for the whole brain over pre-defined periods of interest to assess whether significant MMN and/or P3a emerge (see below). Spatiotemporal clusters of significant differences between standard and deviant ERP amplitudes over each period were identified using non-parametric cluster-based permutation statistics (Blair & Karniski, 1993), implemented with the Fieldtrip toolbox in MATLAB (Maris & Oostenveld, 2007). Using this method, contiguous spatiotemporal clusters of differences between standard and deviant were identified, and the significance of a cluster was determined by comparing the sum of t-values in this cluster to the sum of t-values in the largest cluster identified when standard and deviant labels were randomly permuted for each subject over 10,000 iterations. The associated cluster-level *p*-value was the proportion of random permutations that resulted in a cluster larger than the one identified.

We used cluster-based permutation statistics to identify electrode sites and temporal windows where a significant MMN emerged (defined for our study as a negativity within the 150-300 ms time range) or where a significant P3a emerged (for our study, a positivity in the 300-450 ms time range). These clusters were identified across all participants, separately for each of the four deviant types (harmonic, inharmonic, no-beating, and beating). The difference values (deviant minus standard) within each spatiotemporal cluster were then averaged together for comparison by group (amusic or control), feature (beating or harmonicity) and consonance (consonant or dissonant).

## 3. RESULTS

### 3.1. Elevated discrimination thresholds for harmonicity and beating cues in amusia

Inharmonic shift thresholds were increased for amusic participants relative to control participants [t(1,21) = 2.24, *p* = 0.036, d = 0.94] (see Figure 3, top left). A similar effect was observed for both of the beating-related tasks: complex single-sideband beating detection [t(1,21) = 2.91, *p* = 0.0084, d = 1.23] (Figure 3, top middle) and pure-tone AM detection [t(1,21) = 2.98, p = 0.0071, d = 1.27] (Figure S1). Although the deficits associated with amusia are often considered to be pitch specific, here amusics exhibited reduced behavioral sensitivity to non-pitch cues in beating and AM. The similar effect sizes suggest that the beating deficit may be as important for amusia as the harmonicity deficit. As expected, detection of beating in a complex tone was comparable to detection of simple AM carried by a pure tone. Indeed, thresholds for these two tasks correlated with each other [r = 0.73, *p* < 0.001]. A weaker correlation was observed between AM detection and harmonicity detection [r = 0.56, *p* = 0.005] (Figure S1), with no significant correlation between thresholds for beating and harmonicity [r = 0.30, *p* = 0.16] (Figure 3, top right).

**Figure 3.**
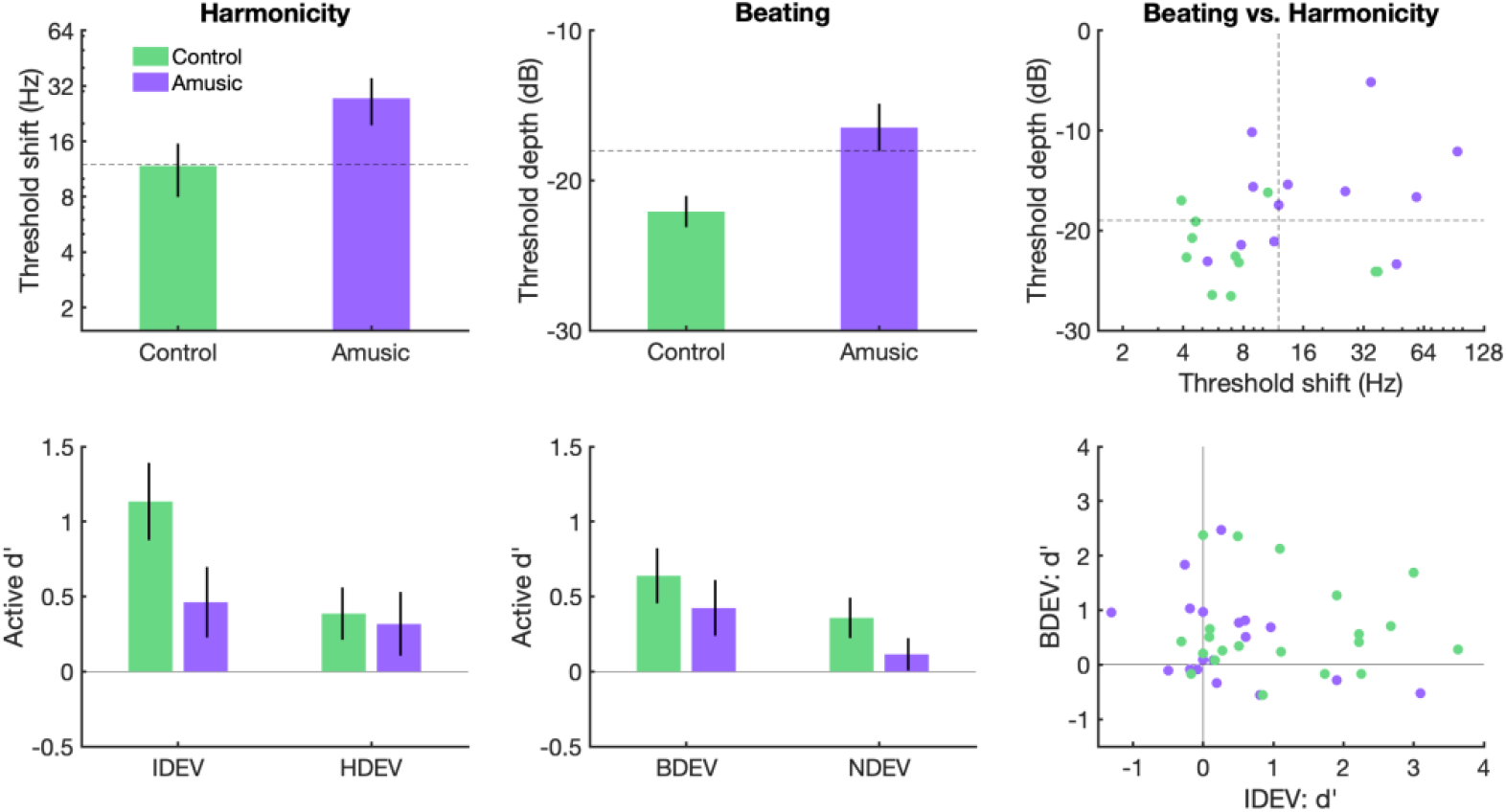
Behavioral results for discrimination of harmonicity and beating cues by amusic and control participants. **Top**: thresholds measured with 3AFC adaptive tracking for inharmonic shift (harmonicity) and single sideband depth (beating), for controls (green, N = 11) and amusics (purple, N = 12). Dashed lines indicate levels chosen for the constant-stimuli oddball paradigm in the EEG study. Left and middle panels: mean per group, error bars show ±1 SEM. Right: individual data. **Bottom**: behavioral sensitivity (d′) for identifying deviants during active task in the EEG study, in the inharmonic deviant (IDEV), harmonic deviant (HDEV), beating deviant (BDEV), and no-beating deviant (NDEV) conditions, for controls (N = 21) and amusics (N = 19). Chance performance corresponds to a d′ of 0.

### 3.2. Decreased behavioral sensitivity to harmonicity and beating changes in amusia

When tested behaviorally during the active task blocks of the EEG experiment, with stimulus differences at a constant level between median control thresholds and median amusic thresholds, the behavioral impairment observed in the threshold measurement study was again confirmed with impaired deviant detection for amusics for both cues. We conducted a nonparametric^b^ version of a 3-way mixed ANOVA on sensitivity (d′) considering the between-subjects factor of participant group (amusic or control) and within-subjects factors of feature (harmonicity or beating) and consonance (consonant or dissonant), using the nparLD package in R (Noguchi et al., 2012). We found a main effect of consonance [F(1,38) = 10.61, *p* = 0.0011], reflecting better performance on dissonant-deviant blocks (IDEV and BDEV) than on consonant-deviant blocks (HDEV and NDEV). We also found a main effect of participant group [F(1,38) = 4.74, *p* = 0.029], reflecting better performance for controls than amusics. No other significant main effects or interactions were found (all *p* > 0.35).

Some participants had previously participated in the threshold measurement study, in which the 3AFC task with its dissonant target resembled the dissonant-deviant conditions more than the consonant-deviant conditions. In order to test whether the apparent effect of consonance was merely an effect of practice from the behavioral threshold study, we re-ran the ANOVA excluding the 13 participants who had completed threshold measurements. The main effect of consonance remained after removing these subjects [F(1,25) = 5.09, *p* = 0.024], suggesting that this effect was not due to participants having practiced, and may reflect an inherent perceptual asymmetry in the stimuli, perhaps due to processes of auditory feature extraction (Cusack & Carlyon, 2003).

Participant performance was evaluated against chance with t-tests of d′ against zero. Control participants performed significantly better than chance in all four conditions (IDEV: *p* < 0.001, HDEV: *p* = 0.04, BDEV: *p* = 0.0024, NDEV: *p* = 0.016), while amusic participants only performed better than chance in the BDEV condition (*p* = 0.046). If Bonferroni multiple-comparisons correction was applied (α = 0.05 / 8 = 0.00625), only control participants ever performed better than chance, and only in the IDEV condition. These results, along with the significant main effect of group on d′, suggest that behavioral discrimination of the stimuli used in the EEG study was more difficult for amusics than for controls, as expected based on the threshold study.

### 3.3 Presence of the MMN and P3a in response to consonant sounds

Each deviant ERP was compared against its acoustically identical standard (see Figure 4). In general, peak negativity of deviants relative to standards occurred in frontal electrodes between 150 and 300 ms, followed by positivity in frontal electrodes between 300 and 450 ms (see Figures 5 and S2). Significant MMNs were observed only for consonant deviants.

**Figure 4.**
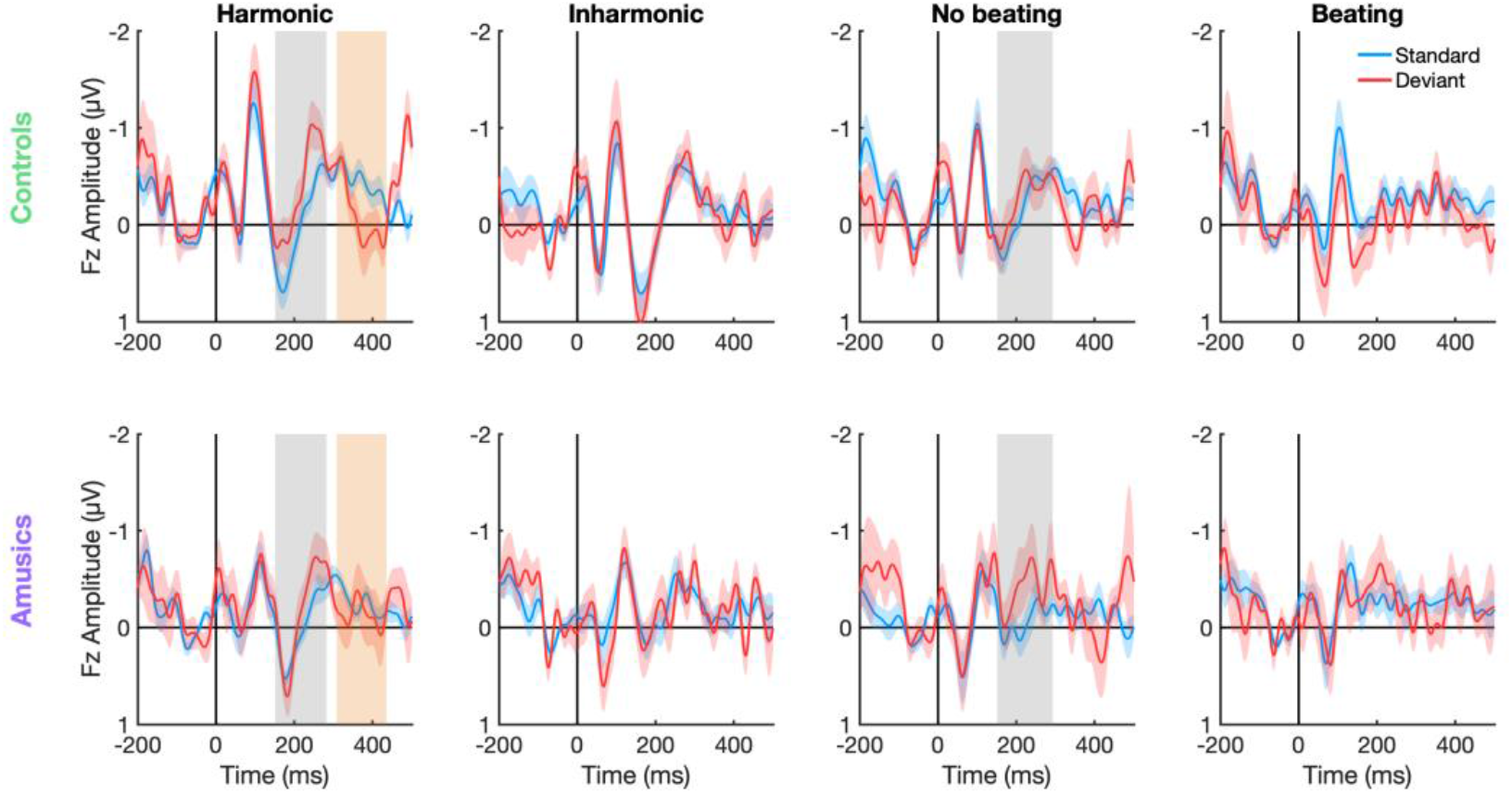
ERPs to standard and deviant stimuli at Fz by group and condition. Responses are shown at electrode Fz for each of the four types of stimulus, for controls (top row) and amusics (bottom row). Each plot compares a deviant to its acoustically identical standard (the same sound in a different context from the opposite condition). Blue and red colored regions show ±1 SEM. Gray regions indicate significant MMN across all participants, observed for harmonic and no-beating sounds. Orange regions indicate significant P3a across all participants, observed for harmonic sounds.

**Figure 5.**
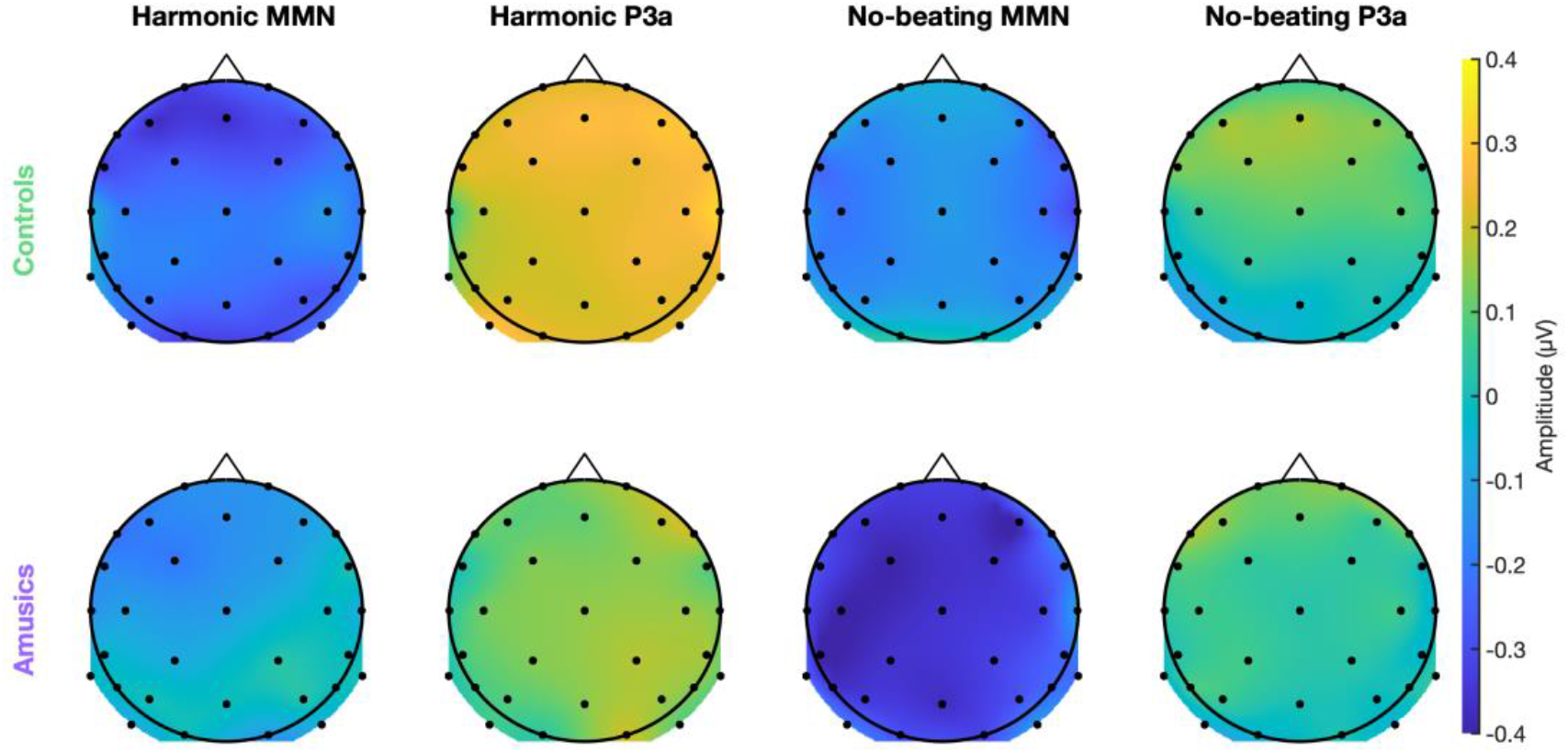
Topography of MMN and P3a responses to consonant stimuli. The difference amplitude (deviant minus standard) across the scalp is shown, averaged over two time periods: 150-300 ms (MMN, odd columns) and 300-450 ms (P3a, even columns), for harmonic sounds (left) and no-beating sounds (right), for controls (top) and amusics (bottom). Small black dots indicate electrode positions. See Figure S2 for the topographies of non-significant responses for dissonant deviants.

Across all participants, significant MMN clusters were identified for harmonic deviants compared to harmonic standards (*p* = 0.034, 150-290 ms) and no-beating deviants compared to no-beating standards (*p* = 0.011, 150-300 ms), but not for inharmonic or beating stimuli (cluster-level *p* > 0.73). See Figure S3 for the temporal course of MMN topographies. A significant P3a cluster was identified for harmonic stimuli (*p* = 0.026, 300-440 ms), but not for the other three sounds (cluster-level *p* > 0.48).

### 3.4 Amplitude of the MMN by consonance, group, and feature

We compared the amplitude of ERPs across participant groups, features, and consonance levels by running a 3-way mixed ANOVA on ERP amplitudes averaged within the relevant spatiotemporal cluster: for harmonic and inharmonic stimuli, the harmonic MMN cluster; for no-beating and beating stimuli, the no-beating MMN cluster (see Figure 6). We observed a main effect of consonance on MMN amplitude [F(1,38) = 15.13, *p* < 0.001, *η*^*2*^ = 0.28], reflecting larger MMNs for harmonic and no-beating stimuli than for inharmonic and beating stimuli, in keeping with the results of the detection tests per condition reported above. This effect ran opposite to the effect of consonance on behavior, as stronger MMNs were observed in response to consonant deviants, but better behavioral performance for dissonant deviants. As with the behavioral results, we re-ran this analysis without the 13 subjects who had completed threshold measurement before the EEG experiment, in order to check whether it might be related to practice on the 3AFC threshold task with dissonant targets. After removing these subjects, the effect of consonance on MMN strength persisted [F(1,25) = 13.18, *p* = 0.0013, *η*^*2*^ = 0.35], suggesting it did not depend on practice in the threshold task.

**Figure 6.**
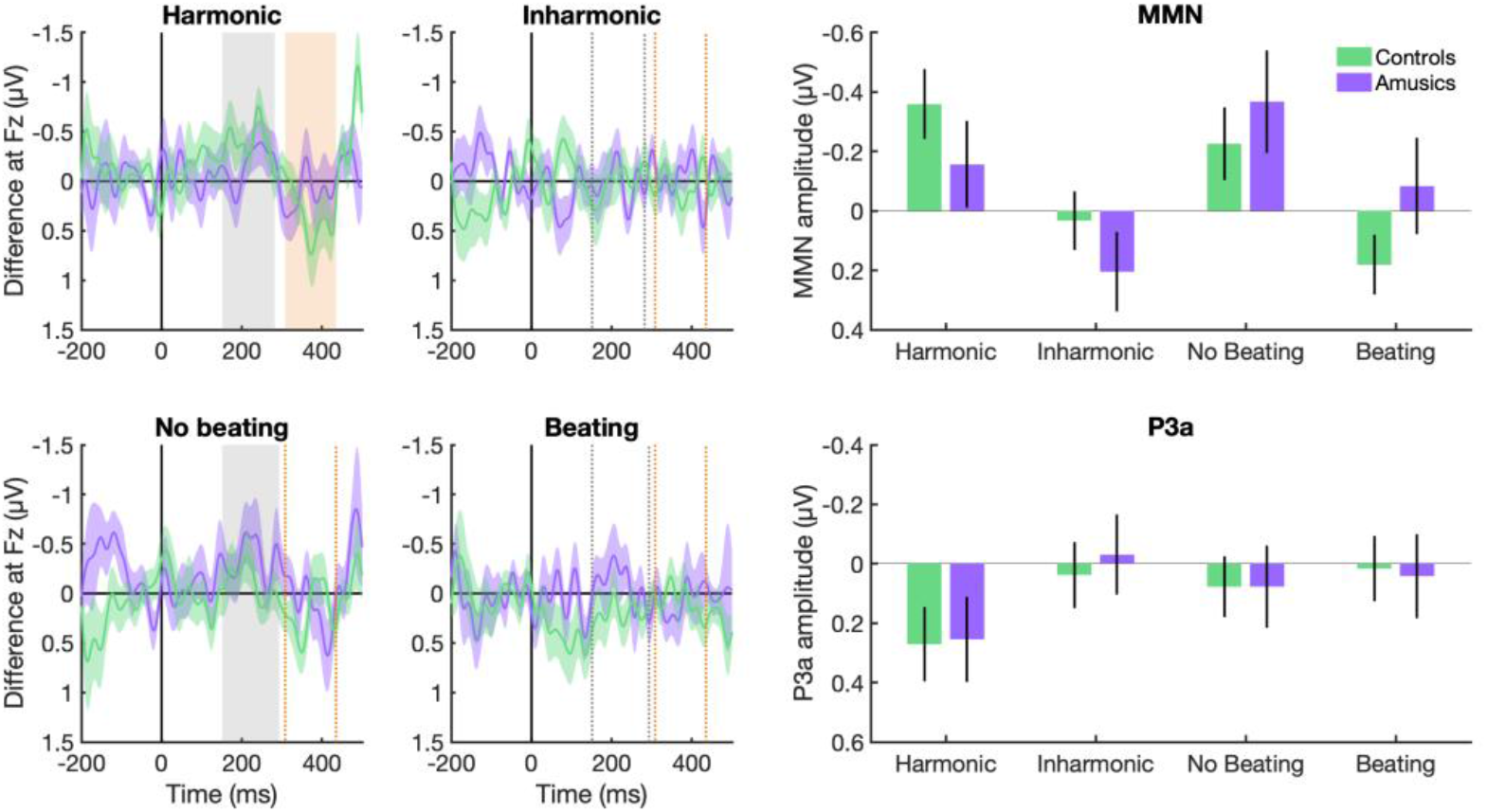
MMN amplitude and difference waves by group and condition. **Left**: Difference waves at Fz for controls (green) and amusics (purple) in each condition. Purple and green colored regions show ±1 SEM. Shaded gray regions show significant MMN at Fz across groups in response to harmonic and no-beating stimuli, while dashed gray rectangles show these same clusters for comparison for inharmonic and beating sounds, where no MMN was observed. The shaded orange region shows a significant P3a for harmonic stimuli, while the dashed orange rectangles show this same cluster for comparison for other sounds where no P3a was observed. **Top right:** mean MMN amplitude, averaged within the relevant spatiotemporal cluster for each group and condition. Error bars show ±1 SEM. For harmonic and no-beating stimuli, averages were computed in the cluster for that stimulus across both groups. As no significant clusters were identified for dissonant stimuli, inharmonic and beating stimuli borrowed the clusters for harmonic and no-beating stimuli, respectively. **Bottom right**: as top right, but for P3a amplitude, with other sounds borrowing the cluster for harmonic stimuli.

We also observed an interaction between group (amusics vs. controls) and feature (beating vs. harmonicity) [F(1,38) = 4.24, *p* = 0.046, *η*^*2*^ = 0.10]. The direction of the interaction (see Figure 6) suggests that MMNs in the control group were larger for harmonicity cues than beating, but MMNs in the amusic group were larger for beating cues than harmonicity. However, post-hoc between-group comparisons (independent t-tests) showed no significant differences between amusics and controls for harmonicity cues [t(1,38) = −1.26, *p* = 0.22] and a marginally significant difference for beating [t(1,38) = 1.78, *p* = 0.084]. Within-group comparisons (paired t-tests) also showed no significant differences between MMN amplitude to harmonicity and beating cues, for amusics [t(1,18) = −1.68, *p* = 0.11] or controls [t(1,20) = 1.17, *p* = 0.25]. The interaction was insignificant when the analysis was restricted to include only consonant-deviant conditions, where significant MMNs were observed [F(1,38) = 1.78, *p* = 0.19].

No other significant main effects or interactions were identified in the ANOVA (all *p* > 0.57). The notable lack of main effect of group means that we found no evidence for impaired MMNs to consonance cues for amusics overall. This is in contrast to our behavioral results, where amusics were impaired overall for both cues, in terms of threshold and sensitivity.

In order to confirm the presence of an MMN for each group in each condition, we compared averaged MMN strengths against zero using one-tailed t-tests, since the expected direction of the effect is negative. Only three MMNs were significant using this test: of controls to harmonic stimuli [t(1,20) = −3.06, *p* = 0.0031], of controls to no-beating stimuli [t(1,20) = −1.86, *p* = 0.039], and of amusics to no-beating stimuli: [t(1,18) = −2.13, *p* = 0.024]. These results, combined with the observed interaction effect, suggest a comparative advantage, in terms of early brain responses, for harmonicity cues for controls and beating cues for amusics.

In order to evaluate P3a amplitudes across group and condition, we ran a 3-way mixed ANOVA on ERP amplitudes in the P3a cluster identified for harmonic stimuli. However, no main effects or interactions were observed (all *p* > 0.12). Limiting the analysis to harmonic stimuli where a P3a response was observed, we observed no significant difference in P3a amplitude between amusics and controls [t(1,38) = 0.09, *p* = 0.93]. In order to verify the presence of P3a response for each group in each condition, we compared averaged P3a strengths against zero using one-tailed t-tests. Only the P3a responses to harmonic stimuli were significant, for both controls [t(1,20) = 2.16, p = 0.022], and amusics [t(1,18) = 1.77, p = 0.047].

## 4. DISCUSSION

Amusic participants in our study showed an impaired ability to discriminate dissonant sounds from consonant sounds, whether using beating or inharmonicity cues. Their MMN responses to the same stimuli were of similar amplitude to those of controls, though they showed a comparatively stronger MMN to beating cues than to inharmonicity cues, relative to controls. These findings have implications for understanding congenital amusia, as well as for the processing of consonance and dissonance overall.

### 4.1 Processing of harmonicity and beating in congenital amusia

We observed that amusic individuals have elevated thresholds and decreased behavioral sensitivity for detection of beating as well as of inharmonicity. Although amusics are behaviorally less sensitive to beating cues as well as harmonicity cues, we found that their MMN responses to these cues were of comparable overall strength to control participants. This finding suggests that initial encoding of these dissonance cues may be intact in amusics despite their poor behavioral performance, pointing to higher-level mechanisms as the source for behavioral impairment. Our results thus agree with previous research suggesting that amusia-related impairments originate in a frontotemporal network connecting auditory areas to higher-order areas involved in working memory and conscious judgments (Albouy et al., 2013, 2015; Hyde et al., 2006, 2007, 2011; Leveque et al., 2016).

Contrary to the conclusions of Cousineau et al. (2012), our findings therefore suggest that congenital amusia involves deficits in the perception of beating as well as the perception of harmonicity. The difference between the previous and present studies likely arises from the fact that the present study tested participants at low levels of dissonance, at and near threshold, as opposed to using high levels of beating that were easily detectable for both groups (as in Cousineau et al., 2012). Our finding of behavioral impairment for beating detection (and closely related AM detection) replicates a recent finding that amusics exhibit deficits for AM coding as well as FM coding (Whiteford & Oxenham, 2017), also using threshold measurements.

Nevertheless, the interaction effect we observed in MMN amplitudes suggests that amusics’ encoding of beating is stronger relative to harmonicity, compared to controls. This may in theory reflect either impaired spectral coding in amusics or enhanced AM encoding in amusics. However, the latter explanation seems highly unlikely given the evidence in the present paper and a previous study (Whiteford & Oxenham, 2017) that amusics are behaviorally impaired for AM detection. Still, despite behavioral deficits, our EEG findings suggest that amusics are, at a minimum, not impaired for early processing of AM in the form of beating cues. It might be argued that the increased emphasis on beating cues in amusics could be a primary, congenital difference in brain function of amusics. But perhaps more likely, it may be a secondary result of impoverished spectral information, in which amusics develop relatively heightened sensitivity to information sources that remain available. This phenomenon may also explain why amusics remain able to use acoustic cues, such as loudness (Graves et al., 2019) or non-spectral cues for prosody (Pralus et al., 2018). Under this explanation, the increased emphasis on beating cues could be compared to some examples of enhanced visual perception for people with profound hearing loss (e.g. Bernstein et al., 2000; Bottari et al., 2011), or to enhanced auditory perception for people with low vision (e.g. Kolarik, Cirstea, Pardhan, & Moore, 2014).

### 4.2 Beating and harmonicity cues in consonance and dissonance perception

Behaviorally, participants in our study more easily detected dissonant deviants in a stream of consonant standards than the reverse. This effect is in line with previous evidence for a behavioral advantage for detecting dissonance (Schellenberg & Trehub, 1994), an effect which has also been observed in infants (Schellenberg & Trehub, 1996). More generally, the advantage for detecting dissonance may be an example of perceptual asymmetry due to feature extraction. In both auditory (e.g. Cusack & Carlyon, 2003; Ruggles & Oxenham, 2014) and visual perception (e.g. Treisman & Gormican, 1988), observers are better able to detect the presence of an extracted feature than its absence. The behavioral advantage for dissonance detection suggests that dissonance cues (beating and inharmonicity) may be extracted as features by the auditory system, whereas their opposites (lack of beating and harmonicity) are only detectable as the absence of a feature.

More surprising than the behavioral effect is the MMN advantage we observed for consonant deviants over dissonant deviants, which runs in the opposite direction to behavioral results. It also runs opposite a recently reported effect of larger MMNs for dissonant deviants than for consonant deviants (Crespo-Bojorque et al., 2018). However, the stimuli in that study were slightly different from those in the present study: whereas the previous study used pairs of simultaneous tones forming consonant or dissonant harmonic intervals, the present study used single complex tones containing isolated low-level dissonance cues. The discrepancy between the present study and Crespo-Bojorque et al. (2018) is perhaps due to the fact that all tones in that study were individually harmonic, although dissonant combinations produced inharmonicity and beating in their interactions. A recent study also found larger MMN amplitudes for harmonic deviants than for inharmonic deviants in amusics and controls (Quiroga-Martinez, Basiński, et al., 2021), providing converging evidence for the asymmetry in processing of harmonic and inharmonic sounds, an asymmetry which can interpreted within a predictive coding framework. An earlier study also supports our finding of a clear MMN to harmonic deviants: Jones (2003) presented rare, deviant harmonic complexes in a stream of mostly inharmonic complexes, in a condition comparable to HDEV in the present study, and observed an MMN. However, as the opposite condition was not tested in that paper, it provides no specific agreement with the present study’s finding for increased MMN to consonant deviants over dissonant deviants. In the present study, the MMN advantage for consonant deviants did not interact with group, suggesting that the asymmetry is present for both amusics and controls. In any case, the direction of the MMN advantage for consonant or dissonant stimuli has no direct bearing on the main findings of the present paper, as an MMN in either case indicates an early brain response indexing detection of the change.

### 4.3 Limitations in measurement of the MMN

The present study focused on stimuli with low levels of dissonance by measuring thresholds and presenting MMN stimuli at constant differences near threshold. Although this allowed us to detect a behavioral impairment for beating perception in amusics that was not detected by Cousineau et al. (2012), it also meant that our analyses of behavioral sensitivity were conducted on data near chance performance, and that our EEG analyses were conducted near the threshold for emergence of MMN. The noise inherent in both of these measurements at such low levels may have impaired our statistical ability to detect differences between groups or conditions, as behavioral performance above chance and the presence of an MMN were not confirmed for every group in every condition. This was especially the case for amusic participants, most of whom had behavioral discrimination thresholds above the inharmonicity and beating levels used in the EEG study. Future studies might further explore this phenomenon by presenting stimuli with larger levels of dissonance in order to observe a more robust MMN response. Another possible reason for small overall MMN amplitudes in the present study was the variability of the standard stimulus, which changed in F0 on every trial. The variability of F0 was necessary in our study to ensure that responses could be interpreted as detection of dissonance, and not as perception of a change in pitch. However, most MMN studies are done using a constant, invariable standard, and the variability in our standard stimulus may have led to decreased MMN amplitude.

### 4.4 Conclusions

Our present findings revealed reduced behavioral sensitivity in amusic participants to two cues that have been implicated in consonance and dissonance perception: beating and inharmonicity. However, our EEG results suggest that inharmonicity and beating are both separately encoded at an early level, as they are differently impacted in congenital amusia, with beating cues given relatively more weight than harmonicity cues in participants with amusia. Taken together, our findings are compatible with the view that inharmonicity and beating each contribute to dissonance perception, but that the relative weight of non-spectral cues may be increased for amusic individuals.

## 5. ACKNOWLEDGMENTS

This work was supported by a grant from the Erasmus Mundus Student Exchange Network in Auditory Cognitive Neuroscience, by LabEx CeLyA (“Centre Lyonnais d’Acoustique”, ANR-10-LABX-0060), LabEx Cortex (“Construction, Function and Cognitive Function and Rehabilitation of the Cortex”, ANR-11-LABX-0042) of Université de Lyon, within the program “Investissements d’avenir” (ANR-11-IDEX-0007) operated by the French National Research Agency (ANR), and by NIH grant R01 DC005216.

---

**Figure S1.**
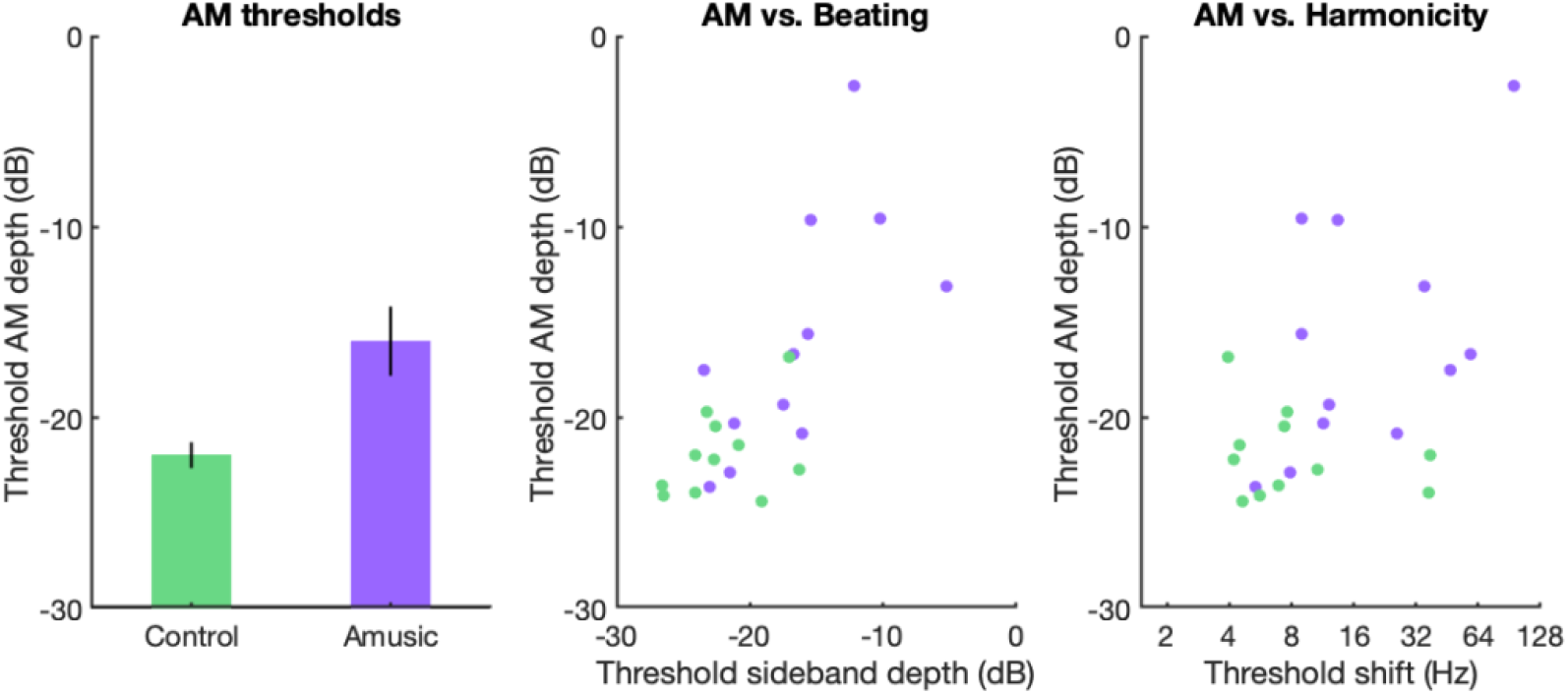
Thresholds for pure-tone AM detection. Thresholds are shown with average and SEM (left), and as individual data compared against single-sideband beating detection thresholds (middle) and against inharmonic shift thresholds (right). Significant correlations were observed between AM and Beating thresholds [r = 0.73, *p* < 0.001] and between AM and Harmonicity thresholds [r = 0.56, *p* = 0.005].

**Figure S2.**
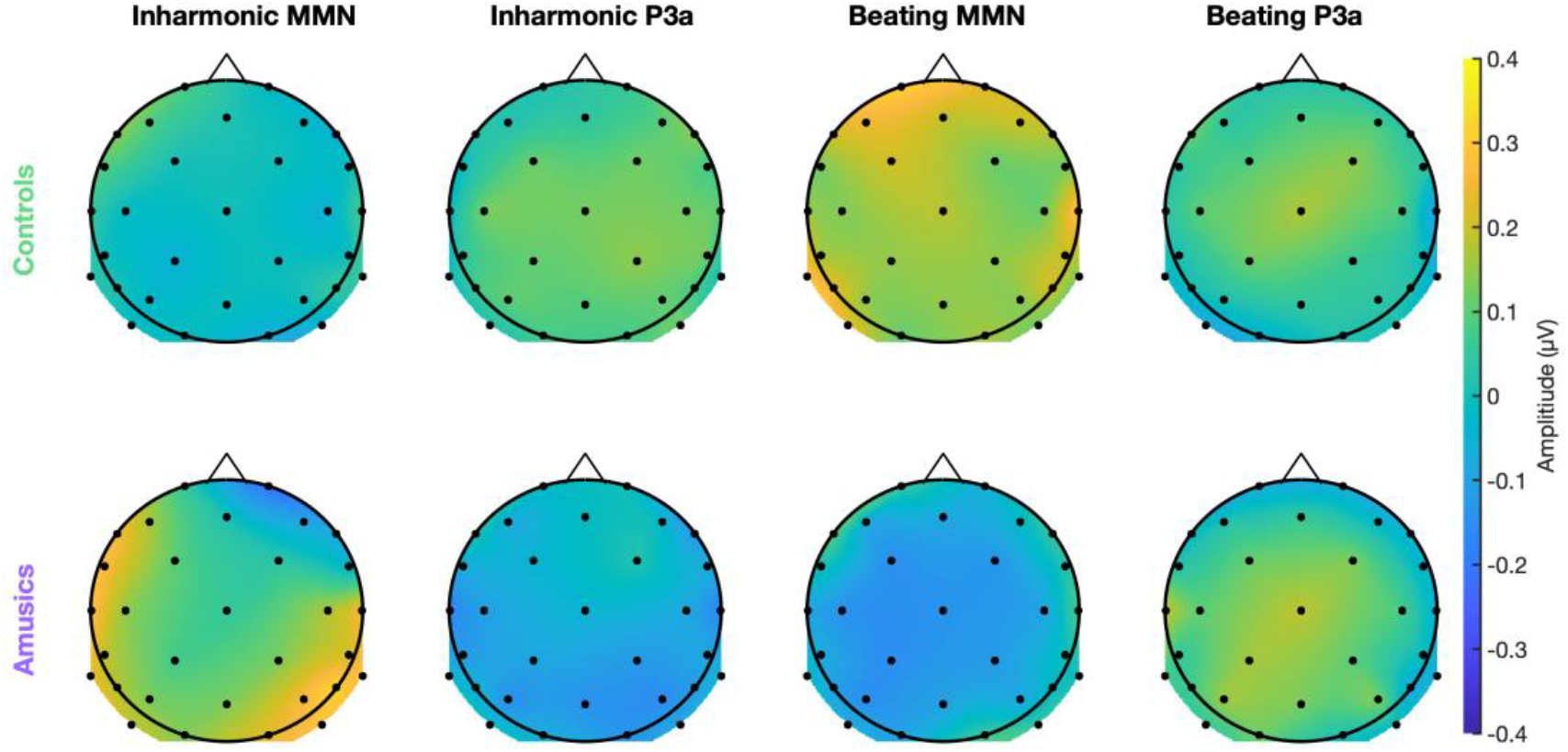
Topography of MMN and P3a responses to dissonant stimuli. As figure 5, but for dissonant stimuli (inharmonic and beating), where no significant MMNs or P3as were observed (as opposed to consonant stimuli shown in figure 5, where significant differences were found).

**Figure S3.**
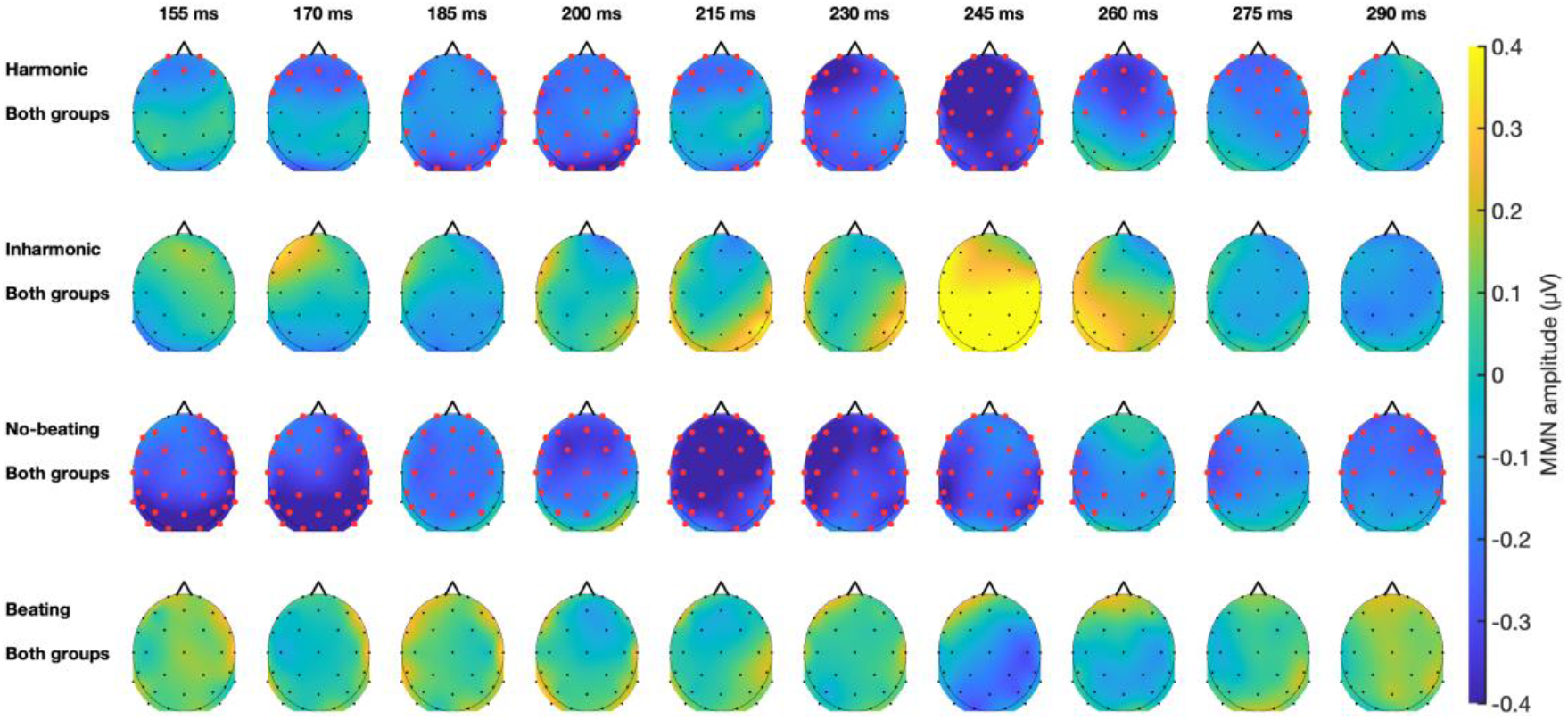
Temporal evolution of MMN topography. The evolution of the difference amplitude (deviant minus standard) for each of four deviant types is shown (rows 1-4), at each of 10 time points (columns 1-10), averaged across all participants. Red dots show electrodes and time points involved in significant MMN clusters for harmonic sounds (row 1) and no-beating sounds (row 3).

## Footnotes

b Because d′ is not normally distributed, with a mode at 0 (representing chance performance), but a longer positive tail (representing better-than-chance performance), a nonparametric test is more appropriate.

## Notes

### Competing Interest Statement

The authors have declared no competing interest.

